# CircRNA hypomethylation in the human amygdala implicates *FKBP5* in alcohol use disorder

**DOI:** 10.1101/2025.05.04.652083

**Authors:** Tara Ghandour, Jill R. Glausier, Arun Asok, Michelle R. Doyle, Paola Campo, Luca Colnaghi, David A. Lewis, Denise B. Kandel, Eric R. Kandel, Giordano de Guglielmo, Shao-shan Carol Huang, Philippe A. Melas

**Author notes:** **Correspondence Philippe A. Melas:** L8:00, Karolinska University Hospital, 17176, Stockholm, Sweden. E-mail address, **Shao-shan Carol Huang:** Center for Genomics & Systems Biology, Department of Biology, 12 Waverly Place, New York, NY 10003, USA. **Author contributions** P.A.M. conceived and designed the study; G.d.G., S.C.H. and P.A.M. supervised research; T.G., M.R.D., and P.C. performed research and/or analyzed data; J.R.G., A.A., L.C., D.A.L., D.B.K., E.R.K., G.d.G., S.C.H., and P.A.M. contributed material, resources, and/or analytic tools; J.R.G., A.A., M.R.D., P.C., L.C., D.A.L., D.B.K., E.R.K., G.d.G., and S.C.H. reviewed and/or edited the paper; T.G., S.C.H., and P.A.M. wrote the paper.

## Abstract

Genome- and phenome-wide association studies implicate the RNA demethylase *FTO* in alcohol use disorder (AUD), yet the RNA methylation landscape in AUD remains poorly characterized. Analyzing postmortem human basolateral amygdala (BLA) tissue, a key brain region in AUD-related behaviors, we found extensive m^6^A hypomethylation uniquely affecting circular RNAs (circRNAs). Notably, *FKBP5*-hosted circRNAs (circFKBP5s) exhibited pronounced hypomethylation correlating with elevated expression of *FKBP5* mRNA isoforms. These findings were replicated in an animal model of alcohol dependence. Predictive analyses suggest that circFKBP5s influence genes involved in neurodevelopmental processes and neuronal identity. These findings uncover a novel aspect of AUD neurobiology linked to circRNA methylation.

## Introduction

Recent genome- and phenome-wide association studies have identified the RNA demethylase *FTO* as a significant genetic risk factor for alcohol use disorder (AUD) (1-3), suggesting that RNA methylation, specifically N6-methyladenosine (m^6^A), may be relevant in AUD pathophysiology. m^6^A methylation is the most abundant internal modification of eukaryotic mRNA, influencing RNA stability, splicing, and translation, with emerging evidence implicating its dysregulation in neuropsychiatric disorders (4). However, AUD-associated changes in mRNA methylation, particularly within brain regions mediating alcohol-related behaviors, such as the basolateral amygdala (BLA) (5), remain largely unexplored. Moreover, little is known about AUD-associated m^6^A modifications on noncoding RNAs, including circular RNAs (circRNAs); a class of stable, brain-enriched RNAs that may contribute to the molecular basis of addiction (6). To fill this knowledge gap, we examined m^6^A modifications across diverse RNA types in postmortem human BLA samples from AUD subjects, and validated key findings in alcohol-dependent rats.

## Results

### Widespread circRNA hypomethylation in the BLA of AUD subjects

We analyzed postmortem basolateral amygdala (BLA) tissue from individuals with alcohol use disorder (AUD) and matched unaffected comparison (UC) subjects (N = 24, **Dataset S1**) using m^6^A methylation arrays targeting multiple RNA types, including mRNAs, circular RNAs (circRNAs), long non-coding RNAs (lncRNAs), and mid-sized non-coding RNAs (e.g., pre-/pri-miRNAs, snRNAs, and snoRNAs). After adjusting for RNA integrity (RIN) and sex, and applying thresholds of FDR < 0.05 and fold change (FC) > 3, we identified 2,635 hypomethylated and 1 hypermethylated circRNA in AUD samples (**Fig. 1A, Dataset S2**). By contrast, we found only a single hypomethylated mRNA (**Dataset S3**), and no significant methylation differences in lncRNAs (**Dataset S4**) or mid-sized non-coding RNAs (**Dataset S5**). To confirm that circRNA hypomethylation was not driven by reduced circRNA expression, we quantified circRNA levels in the same samples using a circRNA expression array. No global reduction in circRNA expression was observed even at lenient thresholds (nominal p < 0.05, FC > 1.5; **Fig. 1B, Dataset S6**). Instead, ten circRNAs were found to be upregulated, including three hosted by the *FKBP5* gene, here termed circFKBP5-1, circFKBP5-2, and circFKBP5-3 (**Fig. 1B;** official circRNA IDs are provided in the legends and *supporting information*)

**Fig. 1.**
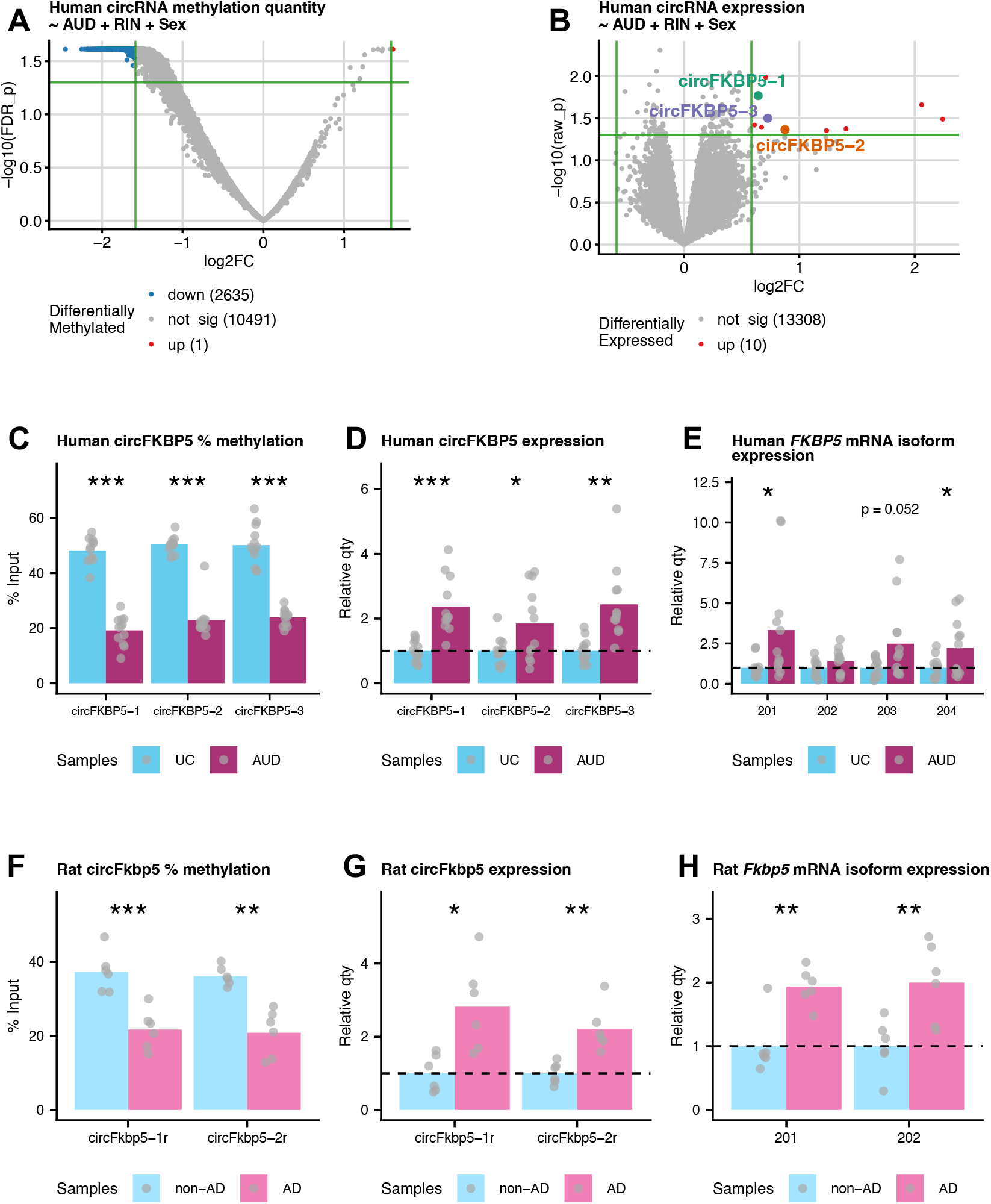
*FKBP5*-hosted circRNAs are hypomethylated and upregulated in the BLA of individuals with alcohol use disorder (AUD) and in alcohol-dependent (AD) rats. ***(A)*** Volcano plot showing differential m^6^A methylation of circRNAs in the BLA of AUD (FDR < 0.05, FC > 3). ***(B)*** Volcano plot showing differential circRNA expression in the same samples (nominal p < 0.05, FC > 1.5); three upregulated circRNAs were hosted by *FKBP5*, termed here circFKBP5-1, -2, and -3. ***(C)*** MeRIP-qPCR validation of circFKBP5 hypomethylation in AUD. ***(D)*** RT-qPCR validation of circFKBP5 upregulation in AUD. ***(E)*** *FKBP5* mRNA isoforms 201 and 204 were significantly upregulated in AUD. ***(F-H)*** Parallel validation in rat BLA: circFkbp5-1r and -2r were hypomethylated ***(F)*** and upregulated ***(G)*** in AD rats, and *Fkbp5* isoforms 201 and 202 were upregulated ***(H)***. Official circRNA IDs in array and/or circAtlas 3.0: circFKBP5-1: hsa_circRNA_076155 (array), hsa-FKBP5_0006 (circAtlas). circFKBP5-2: hsa_circRNA_104101 (array), hsa-FKBP5_0005 (circAtlas). circFKBP5-3: hsa_circRNA_406761 (array), not annotated in circAtlas. circFkbp5-1r: rno-Fkbp5_0003 (circAtlas). circFkbp5-2r: rno-Fkbp5_0006 (circAtlas). Welch’s two-sample t-test: *p < 0.05, **p < 0.01, ***p < 0.001.

### circFKBP5 hypomethylation and expression correlate with *FKBP5* mRNA isoforms

m^6^A methylation has been shown to target certain circRNAs for degradation via the RNase P/MRP complex (7). We therefore hypothesized that the upregulated circFKBP5s in AUD (**Fig. 1B**) may evade degradation due to a hypomethylated state. Supporting this assumption, all three upregulated circFKBP5s were also among the significantly hypomethylated circRNAs in AUD (**Fig. 1A, Dataset S2**). To validate these findings, we used methylated RNA immunoprecipitation (MeRIP)-qPCR, confirming significant hypomethylation of all three circFKBP5s in AUD (p < 0.001, **Fig. 1C**). Follow-up qPCR expression analysis also confirmed their upregulation (p < 0.05, **Fig. 1D**). Given evidence that some circRNAs can enhance expression of their host gene (e.g., by modulating enhancers or promoter methylation) (8), we next examined expression of the four protein-coding *FKBP5* isoforms (201-204 per Ensembl release 113; with 201, 203 and 204 sharing a common transcription start site distinct from isoform 202). Using isoform-specific qPCR, we found that 201 and 204 were significantly upregulated in AUD (p < 0.05, **Fig. 1E**), and isoform 203 showed a trend-level association (p = 0.05, **Fig. 1E**). To test cross-species conservation, we analyzed BLA tissue from alcohol-dependent (AD) and non-dependent (non-AD) rats. Two rat brain-expressed circFkbp5s (here termed circFkbp5-1r and circFkbp5-2r) were both significantly hypomethylated (p < 0.01, **Fig. 1F**) and upregulated (p < 0.05, **Fig. 1G**) in AD rats, mirroring human findings. Furthermore, rat *Fkbp5* isoforms 201 and 202 were also significantly upregulated (p < 0.01, **Fig. 1H**), reinforcing a role for circFKBP5s in regulating *FKBP5* expression.

### Predicted circFKBP5 networks target neurodevelopmental processes

In addition to regulating host gene expression, circRNAs also act in the cytoplasm by interacting with miRNAs and influencing downstream mRNA targets (8). By using overlapping predictions from multiple databases and applying stringent functional filters (see *supporting information*), we identified a human circFKBP5-miRNA-mRNA network involving circFKBP5-1 and circFKBP5-2, which interact with miR-561-5p, miR-708-5p, and miR-642a-5p to target 1,027 mRNAs (990 unique genes; **Fig. 2A, Dataset S7**). Gene ontology (GO) analysis of these targets revealed significant enrichment for neurodevelopment processes, including axon formation and guidance (FDR < 0.001, **Fig. 2B**). To assess cell type specificity of the network, we examined the expression of the predicted mRNAs across 81 human cell types using the Human Protein Atlas single-cell RNA expression dataset (HPA; proteinatlas.org) (9). The strongest enrichment was observed in brain-related cell types, with predicted mRNA targets significantly enriched for excitatory and inhibitory neuron marker genes, supporting the neuronal specificity of the network (FDR < 0.01, **Fig. 2C**).

**Fig. 2.**
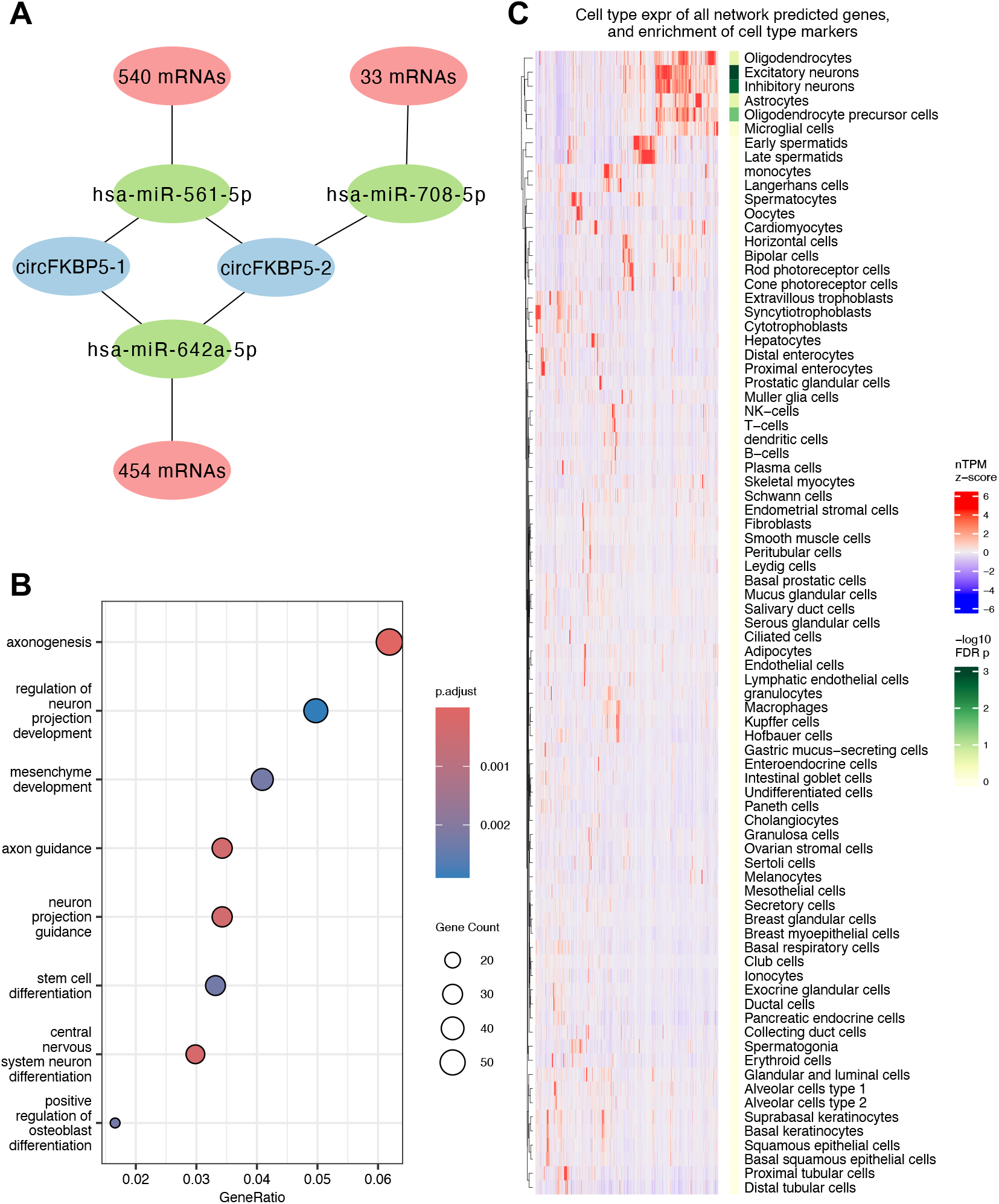
Predicted circFKBP5-miRNA-mRNA network is enriched for neurodevelopmental processes and neuronal marker genes. ***(A)*** Predicted network involving circFKBP5-1 and circFKBP5-2 (both hypomethylated and upregulated in AUD), which interact with miR-561-5p, miR-708-5p, and miR-642a-5p, targeting a total of 1,027 mRNAs (990 unique genes). ***(B)*** GO enrichment analysis of network mRNA targets revealed significant enrichment for neurodevelopmental processes. ***(C)*** Cell-type expression and enrichment analysis using the Human Protein Atlas single-cell RNA expression dataset (9) showed that predicted mRNA targets were most highly expressed in brain-related cell types. Expression values are shown as z-scores of normalized transcript counts (nTPM), with each gene scaled across all 81 normal (non-AUD) human cell types (red = high, blue = low). Green shading between the heatmap and cell type names indicates FDR from enrichment analysis of cell-type marker genes (one-sided Fisher’s exact test).

## Discussion

Our findings reveal a previously uncharacterized epitranscriptomic signature of AUD in the basolateral amygdala (BLA) characterized by widespread m^6^A hypomethylation of circRNAs, a class of brain-enriched transcripts implicated in cognition, memory, and neuronal identity (10-12). We focused on *FKBP5*-hosted circRNAs (circFKBP5s), which were consistently hypomethylated and upregulated in both human and rodent models of AUD. This conserved pattern suggests a potential mechanism in which hypomethylation stabilizes circFKBP5s (7), enabling them to regulate *FKBP5* isoform expression, e.g., through enhancer- or promoter-related mechanisms (8). Given FKBP5’s central role in stress regulation and drinking behaviors (13, 14), and the BLA’s involvement in alcohol-stress interactions (5), our results suggest a novel molecular link between stress response pathways and AUD mediated by circFKBP5 methylation. Beyond transcriptional regulation in the nucleus, circFKBP5s may also act cytoplasmically via miRNA sequestration (8). Our predicted circFKBP5-miRNA-mRNA network identified genes involved in neurodevelopmental processes, particularly axonogenesis and axon guidance. While typically associated with developmental stages, disruptions in these processes may also impact adult brain structure and function, potentially relevant to observed white matter alterations in AUD (15). Although our study establishes correlative relationships, these findings identify circRNA methylation as a promising avenue for future mechanistic research. If causality is confirmed, targeting m^6^A-related pathways, including the activity of demethylases such as FTO, could represent a novel therapeutic strategy for AUD.

### Summary of the Methods

Amygdala samples enriched for basolateral (BL) and lateral (LA) nuclei were obtained from AUD subjects (N = 12; 10 males, 2 females) perfectly matched for sex, and as closely as possible for age and postmortem interval (PMI) to an unaffected comparison subject (UC, N = 12; 10 males, 2 females) through the NIH NeuroBioBank. BLA samples from adult alcohol-dependent rats (AD, N = 6; 3 males, 3 females) and non-AD rats (N = 6; 3 males, 3 females) were obtained from the Alcohol Biobank at the University of California San Diego. m^6^A epitranscriptomic arrays were used to profile RNA methylation, and circRNA expression arrays were used to assess circRNA abundance (Arraystar Inc., Rockville, MD, USA). MeRIP-qPCR and RT-qPCR were conducted to validate RNA methylation and expression levels, respectively. The circFKBP5-miRNA-mRNA network was constructed using functional prediction criteria. Extended methods are provided in the *supporting information*.

### Data, Materials, and Software Availability

All study data and materials/methods are included in the article and/or *supporting information*. Array data are stored in the NeuroBioBank Data Repository (https://nda.nih.gov/nbb) under the collection C5489.

## Acknowledgements

We thank the NIH NeuroBioBank at the University of Pittsburgh School of Medicine that provided human postmortem brain tissue. This work was supported by the Swedish Research Council (Dnr. 2023-02253; P.A.M.), the Swedish Brain Foundation (FO2023-0167 and FO2024-0247; P.A.M.), the Alcohol Research Council of the Swedish Alcohol Retailing Monopoly (FO2023-0046; P.A.M.), the National Institute of General Medical Sciences (R35GM138143; S.C.H.), the National Institute on Alcohol Abuse and Alcoholism (R01AA030048; G.d.G.), and the Howard Hughes Medical Institute (E.R.K.). The funding sources were not involved in the study design or the decision to submit the results for publication.

