## Supporting Information for "CircRNA hypomethylation in the human amygdala implicates *FKBP5* in alcohol use disorder"

### Extended Methods

#### Human subjects and sample characteristics

Brain specimens were obtained by the NIH NeuroBioBank at the University of Pittsburgh during routine autopsies conducted at the Office of Allegheny County of Medical Examiner (Pittsburgh, PA, USA) after consent was obtained from the next-of-kin. An independent team of clinicians made DSM-IV and ICD-10 consensus diagnoses for each alcohol use disorder (AUD) subject using the results of structured interviews with family members and review of medical records. All AUD subjects had a DSM-IV/ICD-10 diagnosis of alcohol dependence (303.90/F10.20) at the time of death. The same approach was used to confirm the absence of psychiatric and neurologic disorders, including alcohol or illicit substance use disorders, in the unaffected comparison (UC) subjects. All clinical diagnoses were made at the NIH NBB "Confirmed" level. All procedures were approved by the University of Pittsburgh Committee for Oversight of Research and Clinical Training Involving Decedents and Institutional Review Board for Biomedical Research. The cohort used in this study consisted of 12 subject pairs (N=24), with each pair composed of one AUD and one UC subject (Dataset S1). To reduce biological variance among groups, subjects in each pair were matched perfectly for sex and as closely as possible for age and postmortem interval (PMI). Subject groups did not significantly differ in mean age (t-test, p=0.80), body mass index (BMI, p=0.06), postmortem interval (PMI, p=0.45), RNA integrity number (RIN, p=0.38), or tissue storage time at -80C (p=0.67). Brain pH did significantly differ (p=0.01) between AUD (6.4±0.2) and UC (6.6±0.2) subjects, consistent with prior findings (1). Dataset S1 provides detailed subject information, including NeuroBioBank IDs, pseudoGUIDs (global unique identifiers), sex, age, RIN, PMI, pH, BMI, and tissue storage times. AUD diagnosis was positively correlated with nicotine ($r = 0.845$, $p = 2.03 \times 10^{-7}$), opioid ($r = 0.642$, $p = 0.0007$), and cocaine dependence/abuse ($r = 0.577$, $p = 0.003$).

#### Rat samples

BLA samples were obtained from heterogeneous stock (HS) rats (2) as part of a study funded by NIAAA (R01AA030048), aimed at conducting a genome-wide association study (GWAS) for alcohol dependence traits and establishing a tissue biobank from fully phenotyped alcohol dependent animals. Adult HS rats (N = 12; 6 alcohol-dependent [AD], 3 males and 3 females; 6 non-dependent [non-AD], 3 males and 3 females) were used. AD rats underwent a chronic intermittent ethanol (CIE) exposure model to induce alcohol dependence (3, 4), involving 14 hours/day of ethanol vapor exposure for 5 weeks. Prior to and following CIE, rats self-administered oral ethanol (0.1 ml, 10% v/v) and water (0.1 ml) on a concurrent fixed-ratio 1 (FR1-FR1) schedule. Phenotypic assessments included alcohol intake (g/kg), escalation index, alcohol preference (alcohol reinforcers/[alcohol + water reinforcers]), motivation via progressive ratio schedule, and compulsive-like drinking using quinine-adulterated ethanol. Ethanol sensitivity and tolerance were evaluated using the loss of righting reflex test, and withdrawal-induced hyperalgesia was measured with the von Frey test. Non-AD rats, serving as age-matched naive controls, did not undergo CIE or ethanol self-administration. At study completion, rats were euthanized during acute withdrawal from alcohol (7-10 hours after last alcohol exposure), and brains were rapidly harvested, snap-frozen, and stored at -80°C. BLA tissue was microdissected using a cryostat to ensure precise anatomical isolation. All procedures were conducted in accordance with Institutional Animal Care and Use Committee (IACUC) guidelines at the University of California San Diego.

**$m^6A$ epitranscriptomic arrays**

To quantify $m^6A$ methylation in BLA samples from individuals with alcohol use disorder (AUD) and unaffected comparison (UC) subjects, we utilized $m^6A$ epitranscriptomic microarrays (Arraystar array service; Arraystar Inc., Rockville, MD, USA). The $m^6A$-circRNA epitranscriptomic microarray (8x15K) captured 13,127 circRNAs (Dataset S2), while the $m^6A$-mRNA&lncRNA epitranscriptomic microarray (8x60K) captured 22,536 mRNAs (Dataset S3), 4,498 lncRNAs (Dataset S4), and 1,514 mid-sized non-coding RNAs (pre-miRNAs, pri-miRNAs, snRNAs, and/or snoRNAs; Dataset S5). Total RNA was extracted and quantified using a NanoDrop ND-1000 spectrophotometer. RNA integrity was assessed via an Agilent 2100 Bioanalyzer or MOPS gel electrophoresis. For each sample, 3-5 µg of total RNA was incubated with an $m^6A$ spike-in control mix in 300 µL of IP buffer (50 mM Tris-HCl, pH 7.4, 150 mM NaCl, 0.1% NP-40, 40 U/µL RNase Inhibitor) and 2 µg anti-$m^6A$ rabbit polyclonal antibody (Synaptic Systems, Cat# 202 003). The mixture was rotated at 4°C for 2 hours. Meanwhile, 20 µL of Dynabeads™ M-280 Sheep Anti-Rabbit IgG (Invitrogen, Cat# 11203D) were blocked with 0.5% BSA, washed, and added to the RNA-antibody mixture to capture methylated RNAs (IP fraction). The remaining unbound RNA was retained as the "Sup" fraction. Binding was continued for 2 hours at 4°C, followed by washes with IP and wash buffers. The IP RNA was eluted using 200 µL elution buffer (10 mM Tris-HCl, pH 7.4, 1 mM EDTA, 0.05% SDS, 40 U Proteinase K, and 1 µL RNase inhibitor) at 50°C for 1 hour, then purified by acid phenol-chloroform extraction and ethanol precipitation. For the circRNA array only, IP and Sup RNAs were treated with RNase R (Epicentre, Inc.) to deplete linear RNAs and enrich circular RNAs. Each IP and Sup RNA sample was spiked with calibration controls, separately amplified and labeled using the Arraystar Super RNA Labeling Kit (AL-SE-005): Sup RNAs were labeled with Cy3, and IP RNAs with Cy5. Synthesized complementary RNAs (cRNAs) were purified (RNeasy Mini Kit, QIAGEN, Cat# 74104), and concentration and dye activity (pmol dye/µg cRNA) were measured using NanoDrop ND-1000. Equal amounts (2.5 µg) of Cy3- and Cy5-labeled cRNAs were mixed, fragmented (with 10× Blocking Agent and 25× Fragmentation Buffer), and hybridized to the arrays in an Agilent hybridization chamber at 65°C for 17 hours. Slides were washed and scanned using an Agilent Scanner G2505C. Scanned images were analyzed using Agilent Feature Extraction software (v11.0.1.1). Raw intensities of IP (Cy5) and Sup (Cy3) signals were normalized using the average $\log_2$ intensity of the spike-in RNAs. Only probes with Present (P) or Marginal (M) QC flags in ≥10 of 24 samples were retained for downstream analyses. The $m^6A$ quantity for each transcript was calculated based on the normalized Cy5-labeled signal from the immunoprecipitated (IP) fraction, as follows:

$$\text{m6A quantity} = \boldsymbol{IP}_{Cy5\ normalized\ intensity}$$

The $m^6A$ quantity is derived by normalizing the raw Cy5-labeled IP signal intensities using the average $log_2$-transformed intensities of spike-in RNAs, as follows:

$$\boldsymbol{IP}_{Cy5\ normalized\ intensity} = \log 2(\boldsymbol{IP}_{Cy5\ raw}) - \text{Average}[\log 2(\boldsymbol{IP}_{spike-in\_Cy5\ raw})]$$

**circRNA expression array**

To quantify circRNA expression levels in BLA samples from AUD and unaffected comparison (UC) subjects, the Arraystar Human circRNA Array V2 platform (8x15K) was used (Arraystar array service; Arraystar Inc., Rockville, MD, USA), which captured 13,318 circRNAs (Dataset S6). RNA concentrations were measured using a NanoDrop ND-1000 spectrophotometer (OD260), and RNA integrity was assessed via denaturing agarose gel electrophoresis. Sample labeling and hybridization followed Arraystar's protocols. Briefly, total RNA was digested with RNase R (Epicentre, Inc.) to degrade linear RNAs and enrich for circular RNAs. The enriched circRNAs were then amplified and transcribed into fluorescent complementary RNA (cRNA) using random priming (Arraystar Super RNA Labeling Kit). Labeled cRNAs were purified with the RNeasy Mini Kit (Qiagen), and their concentration and specific activity (pmol Cy3/µg cRNA) were determined using the NanoDrop ND-1000. For hybridization, 1 µg of Cy3-labeled cRNA was fragmented by adding 5 µL of 10× Blocking Agent and 1 µL of 25× Fragmentation Buffer, followed by incubation at 60°C for 30 minutes. The sample was then diluted with 25 µL of 2× Hybridization Buffer. A total of 50 µL of hybridization solution was applied to the gasket slide and assembled onto the array slide. Slides were incubated at 65°C for 17 hours in an Agilent Hybridization Oven. Following hybridization, the arrays were washed, fixed, and scanned using an Agilent Scanner G2505C. Raw data were extracted using Agilent Feature Extraction software (version 11.0.1.1). Quantile normalization and further data processing were performed in R using the limma package. Low-intensity probes were filtered out, and only circRNAs flagged as "Present" (P) or "Marginal" (M) in at least 10 of the 24 samples were retained for downstream analyses.

**Methylated RNA Immunoprecipitation (MeRIP)**

To validate $m^6A$ circRNA methylation findings in human and rat BLA samples, methylated RNA immunoprecipitation (MeRIP) followed by real-time quantitative PCR (qPCR) was performed (Arraystar MeRIP-qPCR service; Arraystar Inc., Rockville, MD, USA). Official IDs of the analyzed circRNAs (as reported in the human epitranscriptomic arrays and/or circAtlas (version 3.0) for human and rat circRNAs) are listed in the table below. For validation in rats, two brain-expressed circRNAs were selected based on circAtlas (version 3.0) annotations and primer design efficiency. For each MeRIP experiment, 1-3 µg of total RNA was incubated with an $m^6A$ spike-in control and 300 µL of 1× IP buffer (50 mM Tris-HCl, pH 7.4; 150 mM NaCl; 0.1% NP40; 40 U/µL RNase inhibitor) containing 2 µg of affinity-purified anti-$m^6A$ rabbit polyclonal antibody (Synaptic Systems, Cat. No. 202003). The reaction was rotated end-over-end at 4°C for 2 hours. In parallel, 20 µL of Dynabeads™ M-280 Sheep Anti-Rabbit IgG (Invitrogen, Cat. No. 11203D) per sample was blocked with 0.5% BSA at 4°C for 2 hours, washed three times with 1× IP buffer, and added to the RNA-antibody mixture. Immunoprecipitation (IP) was performed with head-over-tail rotation at 4°C for another 2 hours. The supernatant (unbound RNA fraction) was saved separately. Beads were washed three times with 500 µL of 1× IP buffer and twice with

500 µL of Wash Buffer (50 mM Tris-HCl, pH 7.4; 50 mM NaCl; 0.1% NP40; 40 U/µL RNase inhibitor). $m^6A$-enriched RNA was eluted from beads using 200 µL of Elution Buffer (10 mM Tris-HCl, pH 7.4; 1 mM EDTA; 0.05% SDS; 40 U Proteinase K) at 50°C for 1 hour. Both IP and supernatant RNA fractions were extracted using acid phenol-chloroform followed by ethanol precipitation. First-strand cDNA synthesis was performed using SuperScript III Reverse Transcriptase (Invitrogen) according to the manufacturer's instructions. Real-time PCR quantification was performed using the specific circRNA primer pairs listed below. A standard curve was generated using a serial 10-fold dilution series (1 to $10^{-6}$) of cDNA and used to quantify the results. The percentage of input (% Input) for each MeRIP fraction was calculated using the following formula:

$$\%\text{Input}= \frac{2^{-\text{Ct MeRIP}}}{2^{-\text{Ct MeRIP}} + 2^{-\text{Ct Supernatant}}} \times 100\%$$

| circRNA methylation | Primer sequence | Tm (°C) | Length of product (bp) |
|---|---|---|---|
| circFKBP5-1 (human): hsa_circRNA_076155 (array ID), hsa-FKBP5_0006 (circAtlas ID) | Forward:5' CGAAGGAGCAACAGTAGAAAA 3'<br>Reverse:5' TCCCATGCCTTGATGACTT 3' | 60 | 187 |
| circFKBP5-2 (human): hsa_circRNA_104101 (array ID), hsa-FKBP5_0005 (circAtlas ID) | Forward:5' GGAGCAACAGTAGAAAGTTCTCTA 3'<br>Reverse:5' GGCACCTTCATCAGTAGTCATT 3' | 60 | 56 |
| circFKBP5-3 (human): hsa_circRNA_406761 (array ID), not annotated in circAtlas | Forward:5' GAAGGAGAAGACCACGACATT 3'<br>Reverse:5' GCATCAGAAAGGGAACACAG 3' | 60 | 174 |
| circFkbp5-1r (rat): rno-Fkbp5_0003 (circAtlas ID) | Forward:5' AAGGCTACTCAAACCCCAATG3'<br>Reverse:5' CCCTCATCAGTAGTCATTGTCCTT 3' | 60 | 77 |
| circFkbp5-2r (rat): rno-Fkbp5_0006 (circAtlas ID) | Forward:5' TTTAGCCTTGGCAAAGATTGT 3'<br>Reverse:5' AATCATTGGGGTCTCCTCACT3' | 60 | 58 |

**Real-Time qPCR for circRNA and mRNA expression**

To validate the expression levels of circRNAs and *FKBP5*/*Fkbp5* mRNA isoforms in human and rat BLA samples, real-time quantitative PCR (qPCR) experiment were conducted (Arraystar qPCR service; Arraystar Inc., Rockville, MD, USA). Official IDs of the analyzed circRNAs (based on the human expression array and/or circAtlas (version 3.0) annotations for both human and rat circRNAs) are provided in the table below. For rat validations, two brain-expressed circRNAs were selected based on circAtlas (version 3.0) data and efficient primer design. Information on *FKBP5*/*Fkbp5* mRNA isoforms was obtained from Ensembl (release 113). First-strand cDNA synthesis was performed using SuperScript III Reverse Transcriptase (Invitrogen), according to the manufacturer's protocol. Real-time PCR quantification was performed using the specific primer pairs listed below. A standard curve was generated using a serial 10-fold dilution series (1 to $10^{-6}$) of cDNA and used to quantify the results. Expression levels were calculated using either the QuantStudio™ 5 Real-Time PCR System or Rotor-Gene Real-Time Analysis Software

6.0, based on the standard curve. For each sample, the relative expression of target circRNAs or mRNAs was calculated as the ratio of the target gene concentration to that of the housekeeping gene β-actin. For visualization in the figures, expression levels were normalized to the mean of control samples (UC or non-AD), which was set to 1, and values from AUD or AD samples were expressed as fold changes relative to this baseline.

| **circRNA expression** | **Primer sequence** | **Tm (°C)** | **Length of product (bp)** |
|---|---|---|---|
| β-actin (human) | Forward:5’ GTGGCCGAGGACTTTGATTG 3’<br>Reverse:5’ CCTGTAACAACGCATCTCATATT 3’ | 60 | 73 |
| β-actin (rat) | Forward:5’ CGAGTACAACCTTCTTGCAGC3’<br>Reverse:5’ ACCCATACCCACCATCACAC3’ | 60 | 202 |
| circFKBP5-1 (human):<br>hsa_circRNA_076155 (array ID),<br>hsa-FKBP5_0006 (circAtlas ID) | Forward:5’ CGAAGGAGCAACAGTAGAAAA 3’<br>Reverse:5’ TCCCATGCCTTGATGACTT 3’ | 60 | 187 |
| circFKBP5-2 (human):<br>hsa_circRNA_104101 (array ID),<br>hsa-FKBP5_0005 (circAtlas ID) | Forward:5’ GGAGCAACAGTAGAAAGTTCTCTA 3’<br>Reverse:5’ GGCACCTTCATCAGTAGTCATT 3’ | 60 | 56 |
| circFKBP5-3 (human):<br>hsa_circRNA_406761 (array ID),<br>not annotated in circAtlas | Forward:5’ GAAGGAGAAGACCACGACATT 3’<br>Reverse:5’ GCATCAGAAAGGGAACACAG 3’ | 60 | 174 |
| circFkbp5-1r (rat):<br>rno-Fkbp5_0003 (circAtlas ID) | Forward:5’ AAGGCTACTCAAACCCCAATG3’<br>Reverse:5’ CCCTCATCAGTAGTCATTGTCCTT3’ | 60 | 77 |
| circFkbp5-2r (rat):<br>rno-Fkbp5_0006 (circAtlas ID) | Forward:5’ TTTAGCCTTGGCAAAGATTGT3’<br>Reverse:5’ AATCATTGGGGTCTCCTCACT3’ | 60 | 58 |

| **mRNA expression** | **Primer sequence** | **Tm (°C)** | **Length of product (bp)** |
|---|---|---|---|
| β-actin (human) | Forward:5’ GTGGCCGAGGACTTTGATTG 3'<br>Reverse:5’ CCTGTAACAACGCATCTCATATT 3’ | 60 | 73 |
| β-actin (rat) | Forward:5’ CGAGTACAACCTTCTTGCAGC3’<br>Reverse:5’ ACCCATACCCACCATCACAC3’ | 60 | 202 |
| FKBP5-201***** (human) | Forward:5’ GCGGCGACAGGTTCTCTACTTA 3'<br>Reverse:5’ TCTCCAATCATCGGCGTTTC 3’ | 60 | 181 |
| FKBP5-202 (human) | Forward:5’ GAAGGCTAGACTCTCCGACCC 3'<br>Reverse:5’ ACTTTGTCTCCAATCATCGGC3’ | 60 | 246 |
| FKBP5-203 (human) | Forward:5’ GCCCTCCTTCCAGGTTCTCTAC3'<br>Reverse:5’ ATCGGCGTTTCCTCACCATT 3’ | 60 | 175 |
| FKBP5-204 (human) | Forward:5’ GCTCGTTTCTTACATCACCCAC 3'<br>Reverse:5’ CCACAAATACTCCAACCCTCAC 3’ | 60 | 115 |
| Fkbp5-201 (rat) | Forward:5’ AACAACCCCGAGGATAAAGG3’<br>Reverse:5’ TGGTGCCCTCATCAGTAGTCA3’ | 60 | 136 |
| Fkbp5-202 (rat) | Forward:5’ CCAATTAAAAGTTGCTGTTCGG3’<br>Reverse:5’ TCATTGTTACTGGTGCCCTCAT3’ | 60 | 155 |

***Note:** Due to primer design constraints, the FKBP5-201 primer distinguishes FKBP5-201 from isoforms 202 and 203, but not from isoform 204.

**Statistics**

Differential $m^6A$ methylation and circRNA expression analyses from the epitranscriptomic and circRNA expression arrays were conducted using multiple linear regression with the *limma* package in R (5). Expression values were modeled as response variables with AUD status, sex, and RNA Integrity Number (RIN) as covariates. The regression coefficient for AUD was used to calculate differential methylation or expression in AUD relative to UC, adjusted for sex and RIN. For significance, an FDR-adjusted p-value < 0.05 and an absolute fold change (FC) > 3.0 were applied to $m^6A$ methylation data, while a nominal $p < 0.05$ and FC > 1.5 were used for circRNA expression data. For validation experiments using MeRIP-qPCR and RT-qPCR, two-group comparisons were performed using Welch's two-sample t-test, with significance set at $p < 0.05$.

**circFKBP5-miRNA-mRNA network and Gene Ontology**

To construct the human circFKBP5-miRNA-mRNA interaction network, we first identified putative circRNA-miRNA interactions using Arraystar's proprietary prediction software, which integrates algorithms from both TargetScan and miRanda and has been used in previous studies of circRNA-miRNA-mRNA networks (6, 7). The top four predicted miRNA interactions were extracted for the three *FKBP5*-hosted circRNAs that were both hypomethylated and upregulated in AUD (circFKBP5-1 [hsa_circRNA_076155], circFKBP5-2 [hsa_circRNA_104101], and circFKBP5-3 [hsa_circRNA_406761]), yielding a total of 12 unique miRNAs. These miRNA interactions were further validated using the circR web tool (8), which incorporates predictions from miRanda, RNAhybrid, and TargetScan (9-11). In the second step, predicted miRNA-mRNA interactions for the 12 miRNAs were compiled using both the TargetScan (v8.0, September 2021) and miRDB (v6.0, June 2019) databases (12, 13). From miRDB, only functional miRNAs listed in the FuncMir Collection were considered (https://mirdb.org/mining.html). For TargetScan, we included miRNAs categorized as "conserved," "broadly conserved," or "poorly conserved but confidently annotated" (https://www.targetscan.org/cgi-bin/targetscan/vert_80/mirna_families.cgi?db=vert_80&species=Human). Predictions from miRDB (RefSeq IDs) and TargetScan (Ensembl IDs) were merged and converted to HGNC gene symbols using the biomaRt package in R with Ensembl genes version 113 (GRCh38.p14) (14, 15). Only overlapping miRNA-mRNA pairs present in both databases were retained. Further filtering was applied by retaining pairs with miRDB Target Prediction Scores > 50 and TargetScan Cumulative Weighted Context++ Scores (CWCS) < 0, to ensure high-confidence predictions. Based on these criteria, only circFKBP5-1 and circFKBP5-2 were found to form complete circRNA-miRNA-mRNA networks, which were visualized in Cytoscape (16). The predicted mRNA targets from these networks (990 unique genes) were analyzed for functional enrichment using the clusterProfiler package in R (17, 18), focusing on Biological Process Gene Ontology (GO) terms.

**HPA-based cell type mapping of circFKBP5-miRNA-mRNA network genes**

To explore the expression of the network-predicted mRNAs in normal cell types, we analyzed single-cell RNA-seq data from the Human Protein Atlas (HPA; version 23.0, release date 2023.06.19, Ensembl version 109) (19-21). Datasets were downloaded from the HPA "Downloadable data" page (https://www.proteinatlas.org/about/download) under "RNA single cell type data" and "Data from the Human Protein Atlas in tab-separated format." These data compile single-cell transcriptomes from 31 tissues and classify genes into three specificity levels: cell type enriched, group enriched, and cell type enhanced (https://www.proteinatlas.org/about/assays+annotation#singlecell_rna). For enrichment testing, we defined cell type-specific markers as genes annotated as cell type enriched or group enriched, excluding cell type enhanced genes. To ensure consistent gene annotation, all gene symbols were converted to HGNC format based on Ensembl genes version 113 (GRCh38.p14). Of the 990 unique predicted mRNA targets from our circFKBP5 network, 986 were present in the HPA dataset. We visualized the expression of these 986 genes across 81 cell types using a heatmap of z-scored normalized transcripts per million (nTPM), with each gene scaled across all cell types. Adjacent to the heatmap (Fig. 2C), a bar column displays FDR-adjusted p-values from one-sided Fisher's exact tests that evaluated enrichment of cell type-specific markers in our gene set. These tests were based on 2×2 contingency tables comparing whether the genes were included in our network, and whether they were annotated as cell type-specific markers, using all protein-coding genes from Ensembl v113 as the background. Multiple testing correction was performed using the False Discovery Rate (FDR) method.
